# PEO: Plant Expression Omnibus - a comparative transcriptomic database for 103 Archaeplastida

**DOI:** 10.1101/2023.06.22.546049

**Authors:** Eugene Koh, William Goh, Irene Julca, Erielle Villanueva, Marek Mutwil

**Affiliations:** School of Biological Sciences, Nanyang Technological University, 60 Nanyang Drive, Singapore, 637551, Singapore

**Author notes:** Corresponding author: Marek Mutwil, School of Biological Sciences, Nanyang Technological University,60 Nanyang Drive, 637551, Singapore, Singapore.

## Abstract

The Plant Expression Omnibus (PEO) is a web application that provides biologists with access to gene expression insights across over 100 plant species, ∼60,000 manually annotated RNA-seq samples, and more than four million genes. The tool allows users to explore the expression patterns of genes across different organs, identify organ-specific genes, and discover top co-expressed genes for any gene of interest. PEO also provides functional annotations for each gene, allowing for the identification of genetic modules and pathways. PEO is designed to facilitate comparative kingdom-wide gene expression analysis and provide a valuable resource for plant biology research. We provide two case studies to demonstrate the utility of PEO in identifying candidate genes in pollen coat biosynthesis and investigating the biosynthetic pathway components of capsaicin in *Capsicum annuum*. The database is freely available at https://expression.plant.tools/.

## Introduction

The earliest members of the plant kingdom evolved circa one billion years ago from photosynthetic algae. The archetypical characteristics we associate with plants, such as seeds, roots, leaves, flowers, and wood, were evolutionary innovations that allowed plants to colonize diverse habitats successfully. Today, plants form an indispensable component of life on Earth as the primary food producer in the terrestrial ecosystem. Plants being mostly sessile, have also had to evolve means of adapting quickly to their local environments. These adaptations came in the form of complex regulatory and metabolic pathways, which we are now only beginning to unravel.

Studying plant biology and genetics will allow us to improve desirable traits and engineer crops that provide higher yield and/or are more tolerant to various environmental stresses (Niazian, 2019; Nowicka et al., 2018). The use of plants as a source of various medicinal components is also exemplified in the extraction of the analgesic salicins from willow bark or the antimalarial artemisinin from *Artemisia annua* (Desborough and Keeling, 2017; Krishna et al., 2008). The elucidation of biosynthetic pathways of these medicinal compounds could then be harnessed for large-scale production and modification in recombinant systems (Cravens et al., 2019; Paddon et al., 2013; Sabzehzari et al., 2020). Currently, the study of plant gene function requires the characterization of individual genes and their interactors and regulators, which are typically limited to a handful of model plants. Consequently, bioinformatic approaches that integrate genomic, transcriptomic, metabolomic, and phenotypic data have been developed to accelerate the process of gene function prediction (Hansen et al., 2018; Radivojac et al., 2013; Rhee and Mutwil, 2014).

Studying gene co-expression can effectively address this, given the “guilt-by-association” principle, where genes with similar expression are likely to have similar functions (Rhee and Mutwil, 2014). Co-expression analysis allows one to transfer functional annotation to unannotated genes, predict undiscovered members of metabolic pathways, and identify regulator-target relationships (Rao and Dixon, 2019). Plant organs have signature gene expression patterns and similar profiles between organs are correlated to the similarity in function of the organs (Ma et al., 2005). Moreover, many genes exhibiting organ-specific expression are found to be conserved across land plants (Julca et al., 2021). Consequently, a gene’s expression profile across organs can provide clues to a gene’s function. For example, two genes that show similar gene expression profiles across different organs and tissues are likely to be part of the same protein complex (Persson et al., 2005) or biosynthetic pathway (Delli-Ponti et al., 2021) or to be targeted to the same subcellular compartment (Ryngajllo et al., 2011).

Over the last decade, sequencing cost has been falling faster than Moore’s law (Wetterstrand, 2013), giving rise to immense volumes of plant genomic and transcriptomic data. Many online databases with diverse functionalities and plant species have emerged to provide analytical and visualization tools for the scientific community (Lim et al., 2022; Liu et al., 2023; Robinson et al., 2018; Yu et al., 2022). However, no existing database has been designed to cover well-annotated gene expression data. We believe that the full value of RNA-seq repositories is only realized when insights are accessible to biologists working on the genes, including non-model species. We created a user-friendly tool to help non-programmer biologists extract insights about their gene of interest from big data (Tan et al., 2020). A web application to access gene expression knowledge on any gene would be needed to provide an effortless and instantaneous experience for biologists, which requires large volumes of processed data. While the data is publicly available, the interfaces have not been specifically designed for bulk data acquisition for whole kingdoms. We addressed this by designing a parallel-processing pipeline to automatically discover, download and process RNA-seq data for the whole plant kingdom in less than a month (Goh and Mutwil, 2021), making the provision and maintenance of a kingdom-wide resource feasible.

Here, we developed the Plant Gene Expression Omnibus (PEO, https://expression.plant.tools), a web application to provide an interface for biologists to access gene expression insights across >100 plant species. Specifically, the tool addresses how a gene is expressed across different organs, whether it displays organ-specific expression patterns and identifies top organ-specific genes for each organ. The tool also automatically provides a list of the highest correlated genes based on the Pearson Correlation Coefficient for any gene of interest, negating the need for separate analysis. In addition, Pfam and Mapman annotations have also been linked to each gene, allowing for functional searches to identify genetic modules or pathways. The software design considers scalability, with the ability to include more species, other expression conditions, and other gene annotations. The project lays the architectural foundations for a kingdom-wide gene expression analysis resource.

## Methods

### Data source

Supplemental Table S1 details the sources of raw files used and contributors. These were then reorganized into the upload format described in Supplemental Table S2, using custom Python scripts. A total of 103 species were included. The complete list is available in Supplemental Dataset S1. Gene expression matrices were obtained from a previous study from public databases via our pipeline using recommended thresholds of log_10_-normalized number of processed reads and percentage of reads pseudo-aligned to the reference CDS (Goh and Mutwil, 2021), which removed samples with low number and percentage of mapping reads. Gene expression was quantified as transcript per million (TPM) via pseudoalignment to reference coding sequences (CDS) of each species, using kallisto (Bray et al., 2016). Organ annotations were done by manually mapping samples to Plant Ontology (PO) terms, which are precise and consistent references to plant anatomical entities across species (Walls et al., 2019). The expression data and annotations are available on 10.6084/m9.figshare.24257125. Genes were functionally annotated with Mapman annotations via Mercator (Schwacke et al., 2019) for four representative species and were assigned to the remaining species via sequence similarity score from diamond (Buchfink et al., 2015). Mapman was chosen over Gene Ontology (GO) as it is specific to plant pathways and processes (Klie and Nikoloski, 2012). Genes were also annotated with protein families (Pfam) (Mistry et al., 2021). Gene identifiers were standardized to the versions in TPM matrices, and alternative isoforms were excluded. Co-expression analysis was performed using Pearson Correlation Coefficient (PCC).

### Database schema and software architecture

The schema defines how data is modeled and stored in our document-oriented database and can be described with an entity relational diagram (ERD) (Supplemental Figure S1). The chosen database technology is MongoDB, a document-oriented database allowing flexible data modeling to optimize our specific read patterns. This is more suitable than relational databases that enforce data schema normalization, which would incur expensive join operations on every read. The database is hosted serverless on MongoDB Atlas, on AWS.

## Results and discussion

### Querying the Plant Expression Omnibus (PEO)

PEO is an interlinked database that includes 103 species, 4.6 million genes, 5,760 Mapman terms, 7,566 Pfam terms and 59,856 quality-controlled samples, of which 41,592 are annotated with 78 PO terms. To our knowledge, no species-wide gene expression database exists for many of these 103 species. The distribution of data illustrating species composition and organ specificity are presented as a treemap and heatmap, respectively (Figure 1). Not surprisingly, leaves were the most sampled plant organs (97 species contain leaf samples), followed by roots (81), flowers (70), stems (67) and seeds (49)(Figure 1A). The less represented samples represent structures not found in all flowering plants (strobilus, tuber, protonema, spore) or samples that are difficult to collect (trichome, zygote, egg cell). *Arabidopsis thaliana* had the highest number of RNA-seq samples (Ara.th, 72,732), followed by maize (35,309), rice (25,090), wheat (14,137) and other crops (Figure 1B), with the next model organism *Brachypodium distachyon* on the 17^th^ place with 3,004 samples. The annotation depth of Mapman, Pfam and Plant Ontology annotations are also provided (Supplemental Figure S2). The data are sorted by several modes, such as species type, organ specificity, as well as Mapman and Pfam annotation. Such an arrangement allows the user to query for gene expression in specific organs of interest or look for orthologous genes in non-model species. Sequence-based BLAST and annotation-based Pfam searches can be used to identify homologous and orthologous genes of interest across the database. Together with Mapman annotation and organ specificity predictions, putative functions can be ascribed to gene orthologues. Finally, PEO also provides a list of top co-expressed genes for any gene of interest for all species present in the database, allowing for the discovery of specific metabolic or developmental pathways that are spatiotemporally regulated.

**Figure 1:**
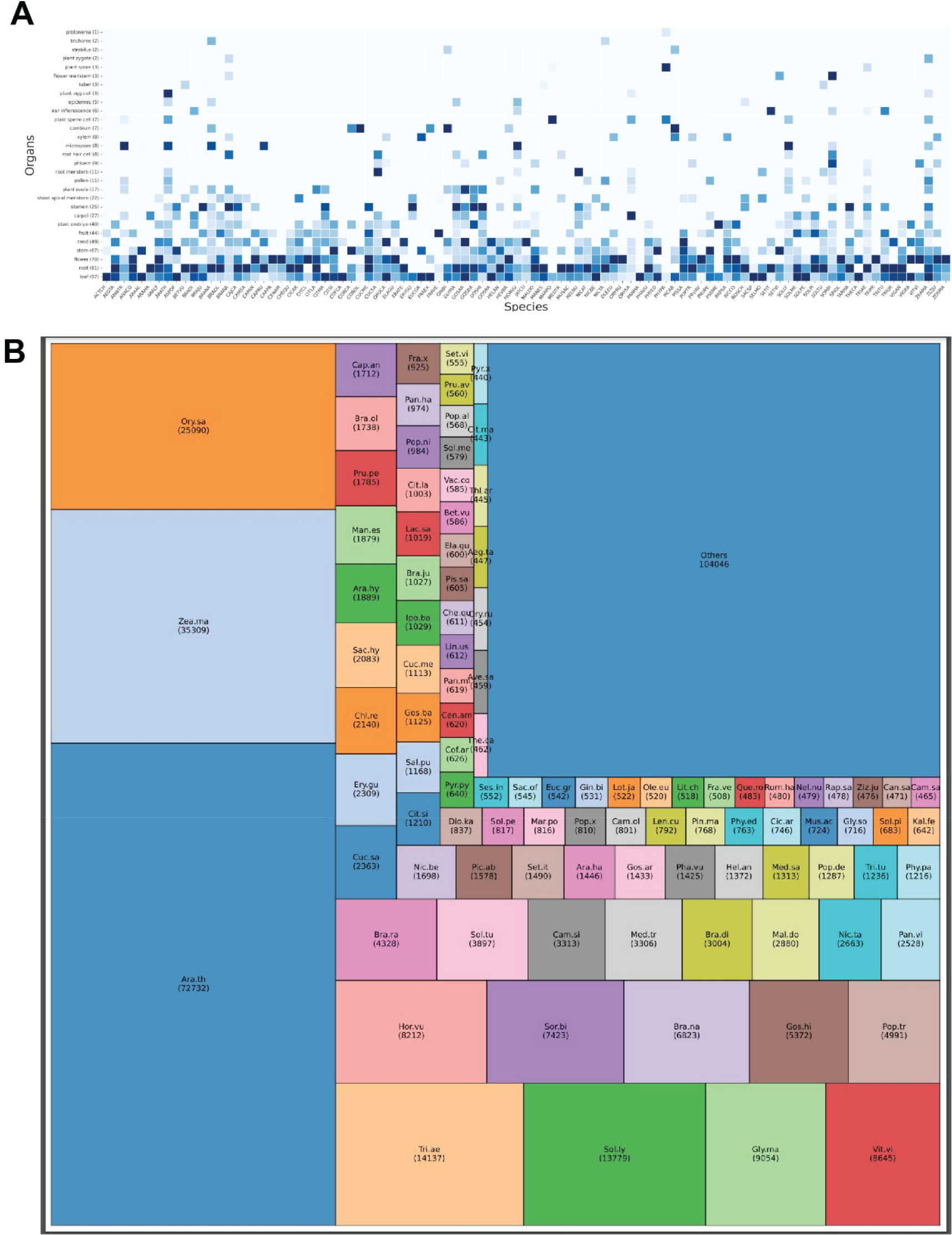
RNA sequencing contents of the Plant Expression Omnibus (PEO) database. A) The heatmap shows the relative abundance of RNA-seq data for the different species (columns) and organs, tissues, and cell types (rows). The numbers in paratheses in rows indicate the number of species that have data for a given plant structure. The cell opacity indicates the relative abundance of RNA-seq samples across organs for each species. B) The treemap showing the relative number of RNA-seq samples for the different species. The areas of the rectangles correspond to the number of samples, which is given in parentheses.

The home page of PEO consists of several search modes. Queries can be based on specific GeneID or protein sequences. It is also possible to browse the database via the ‘Species’, ‘Organs’, ‘Pfam’, or ‘Mapman’ annotations (Figure 2). For example, we can select the ‘Organs’ tab, which provides a list of Plant Ontology (PO) terms. By selecting ‘Pollen’ as our organ of interest, we arrive at a page displaying a set of transcripts sorted by SPM. This specificity measurement ranges between zero and one, where one indicates the gene is exclusively expressed in the tissue (Julca et al., 2021). The page also provides information on the Median and Mean Transcripts per Million (TPM) values used to calculate SPM value and the number of samples the calculations are based on. Mapman annotations are also listed for individual transcripts to provide information on putative gene functions. This database can also be searched by species to obtain the pollen-specific genes in other species present in the PEO.

**Figure 2:**
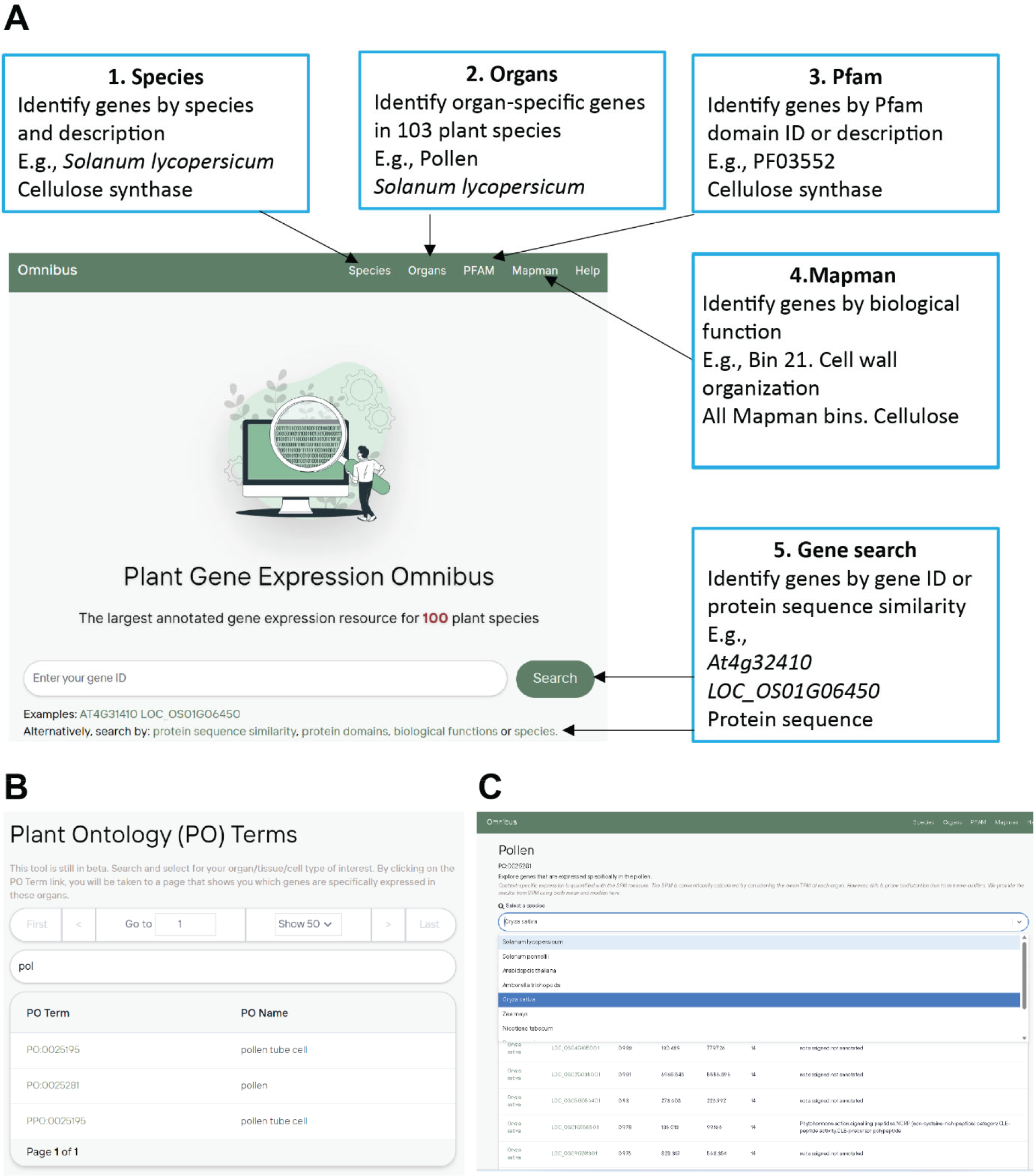
Frontend of Gene Expression Omnibus. A) Genes of interest are searchable via GeneID, protein sequence, Pfam domains, and species as query entries. Browsing of various databases is available in the above tabs. B) Example search by organ type (Pollen). C) The output displays a list of genes known to be present in a specific organ of interest, searchable by species.

As another example of a query strategy, we also show an example search using the protein sequence of the Arabidopsis gene ABORTED MICROSPORES (*AT2G16910*), a transcription factor known to be associated with the regulation of pollen formation (Xu et al., 2014). The search output provides a list of genes sorted by %identity, with further information on E-values, bit scores, number of mismatches, and gap openings. The gene list includes matched genes from the other species present in PEO, providing a means to discover gene orthologues from less common organisms (Supplemental Figure S3).

Clicking on a gene of interest in the list (AMS, *AT2G16910*), brings us to a gene page containing relevant information on this gene (Figure 3). The top of the page provides information about the GeneID, Species name, and TaxonID. The next section displays the putative organ specificity of the gene of interest based on SPM calculations. Details on the values that the SPM calculation is based on is present in the grey subtext below the SPM value. The top organs are reported with mean TPM, standard deviation, and median TPM, showing the magnitude of expression. Since genes with low TPM values can appear to be specifically expressed due to noisy expression, the mean TPM values can be used to flag such genes (Proost and Mutwil, 2018). The sample size is also reported. Together, these support user interpretation of whether the gene displays organ-specific expression. The data can be visualized in a bar chart or box/scatterplot format.

**Figure 3:**
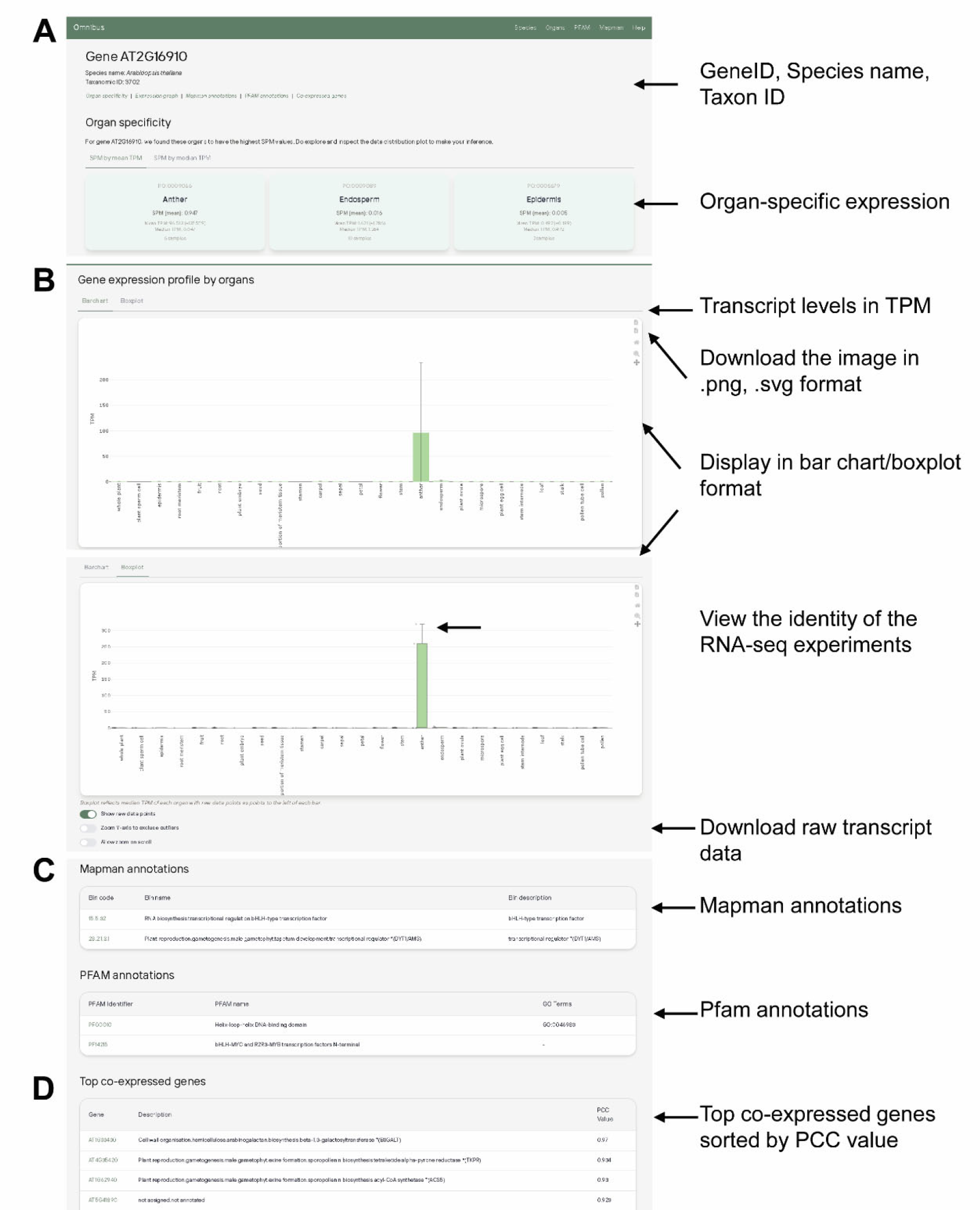
Example gene page of the gene of interest (*AT2G16910*). A) The gene page provides organ specificity of the gene predicted by the SPM score. B) It also displays relative transcript expression in various plant organs/tissues as a box plot or bar chart. Moving the mouse pointer over the points in the box plot shows the corresponding experiment ID. C) Mapman and Pfam annotations for the gene of interest are displayed to provide information on gene function. D) A list of top co-expressed genes sorted by PCC value.

The boxplots depict quartiles and whiskers (hover to reveal values), indicating the skew of data. PEO allows for automatic rescaling to exclude the outliers for the y-axis (Figure 3). To see which experiments the data stems from, the user can move the mouse pointer over individual data points, which will reveal the run IDs. Manual zooming and panning is also available by the control bar at the right of each chart. The charts are available to download in .png or .svg image formats, with the raw data for the annotated samples also available for download in tsv format. Associated Pfam and Mapman annotations are also provided, with the relevant Mapman bin codes and Pfam names and identifiers available for reference. Lastly, a list of top-coexpressed genes sorted by Pearson Correlation Coefficient (PCC) is provided for the user.

### Use case 1: Identification of genes involved in pollen coat biosynthesis

Here, we provide a case study on how PEO can be used to identify candidate genes in pollen coat biosynthesis by using PEO and correlation with other evidence available in the literature. As a starting point, we will use *ABORTED MICROSPORES* (*AMS, AT2G16910*), a transcription factor known to regulate pollen coat biosynthesis (Xu et al., 2014). The pollen coat is a multilayered structure that provides protection for the nascent male gametophytes from external stresses (Scott et al., 2004). The outer layer, called exine, is composed primarily of a polymer called sporopollenin, which is a composite of fatty acids, phenylpropanoids, phenolics, and carotenoids (Ahlers et al., 1999). It was found that the *ams* null mutant failed to produce functional pollen, and further investigations led to its identification as a master regulator of various aspects of pollen biosynthesis (Xu et al., 2014).

Using PEO, we entered its GeneID (AT2G16910) in the search bar on the home page. We first looked at the gene page of AMS, which showed its highly specific expression in the anthers (Figure 4). Next, we obtained the list of top co-expressed genes against *AMS* and used it to compare against gene clusters identified by FlowerNet. FlowerNet is a flower-specific database based on microarray data focusing on stamen-, pollen-, or flower-specific expression (Pearce et al., 2015). By generating a correlation network, FlowerNet generates gene clusters of transcriptionally coregulated genes, and based on analysis of tissue specificity and expression at different pollen maturation stages, we were able to identify several clusters associated with pollen development. Venn diagram analysis showed that AMS co-expressed genes overlapped significantly (55%, p = 1.4E-26) with Cluster 37, representing sporopollenin biosynthesis, but not Clusters 21 and 116, representing pollen exine formation and pollen tube growth, respectively (Figure 4A). This suggests that *AMS* plays a key role in regulating sporopollenin biosynthesis in pollen development.

**Figure 4:**
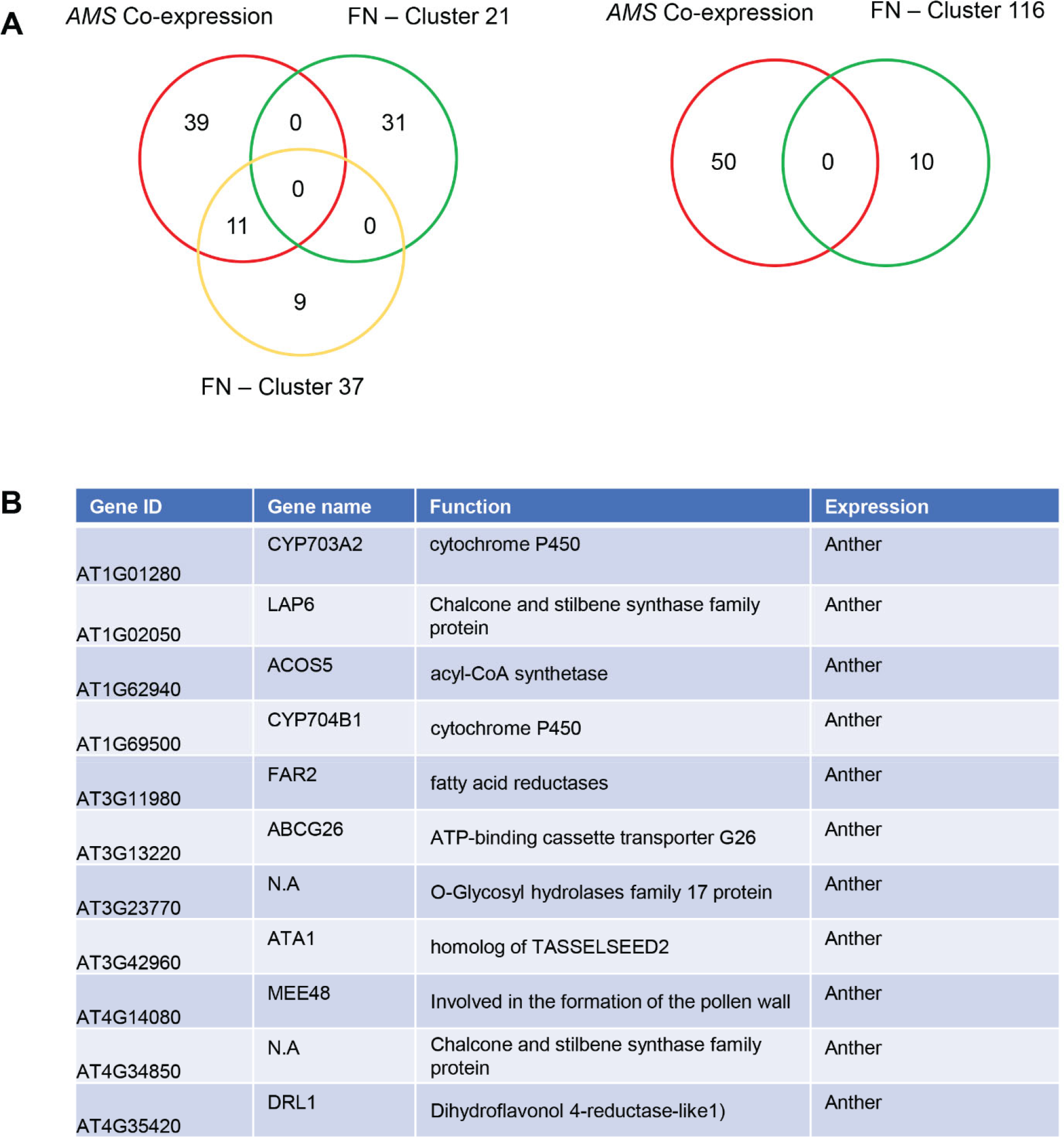
Example use case of gene *AMS* (*AT2G16910*), a transcription factor known to regulate pollen coat biosynthesis. A) Gene expression profile by organs shows its highly specific expression in pollen. Comparison of top co-expressed genes with co-expression clusters extracted from FlowerNet shows strong overlap with Cluster 37 (Sporopollenin biosynthesis) but not Cluster 21 (Pollen exine formation) and Cluster 116 (Pollen tube formation/growth). B) Table shows gene names, functions, and specific expression of genes found in the overlap between AMS co-expression and FlowerNet Cluster 37.

Further examination of the overlapped genes (11 genes) revealed a list of genes that were annotated as being involved in pollen development (*AT1G01280, AT1G69500, AT4G35420*) or whose mutants were essential for male fertility (*AT1G62940, AT3G11980, AT3G13220, AT4G35420*), and were all found to be predominantly expressed in anthers (Figure 4B, Supplemental Dataset S2). This strategy of using complementary databases can be used to narrow down large gene lists and identify key genes involved in a pathway of interest. Finally, using the associated Mapman and Pfam annotations available on the gene page, we can obtain gene orthologues from other species and use that as a starting point to investigate genetic pathways in other plants (Figure 5A). As an example, we examined the genes *SOLYC02G079810*.*3*.*1* and *LOC_OS07G36460*.*1* from *Solanum lycopersicum* and *Oryza sativa*, respectively (Figure 5B). Observations of the respective gene loci of *Solanum lycopersicum* and *Oryza sativa* show transcript enrichment in reproductive tissues similar to that in Arabidopsis.

**Figure 5:**
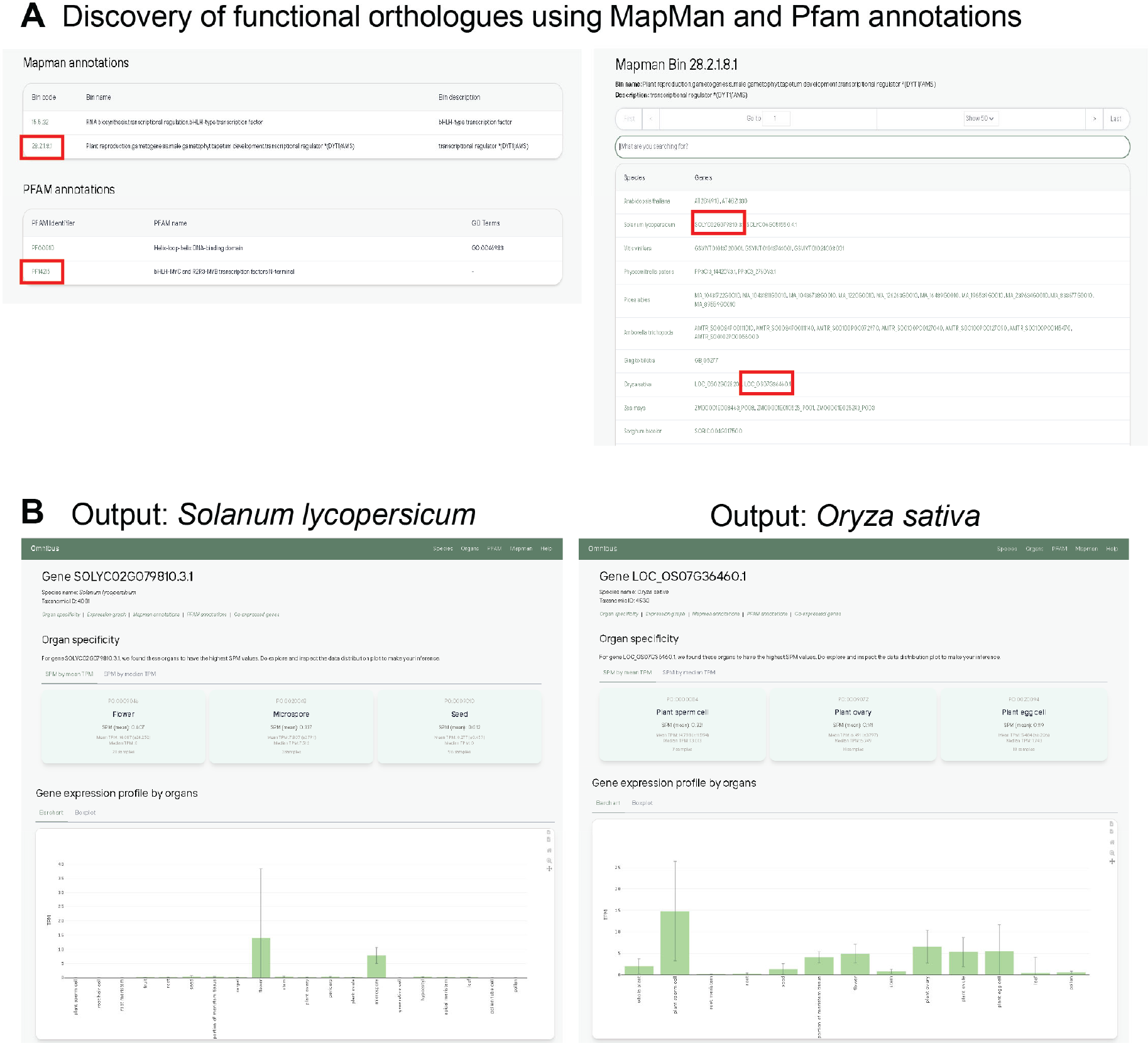
Discovery of functional orthologues using MapMan and Pfam annotations. A) Mapman and Pfam annotations for the gene of interest (*AT2G16910*) provide a list of genes annotated with the bins and domains in other species. B) Example output for functional orthologue discovery using MapMan annotation. The gene of *Solanum lycopersicum* and *Oryza sativa* show specific expression in reproductive tissues similar to *AT2G16910* in *Arabidopsis*.

### Use case 2: Identification of biosynthetic pathway components of capsaicin

Here, we will provide an example of using PEO to investigate capsaicin biosynthesis in the crop plant *Capsicum annuum*. The genus Capsicum comprises a large class of crop plants commonly known as peppers (i.e. chili peppers), and its distinctive pungency/spiciness is due to the alkaloid capsaicin (Suzuki et al., 1980). We looked at the literature and found that *Pun1* (*CA02G19260*) is a necessary enzymatic component of capsaicin biosynthesis. The *Pun1* gene encodes an acyltransferase, and the deletion of the *Pun1* gene in bell peppers is the reason for its distinctive non-pungency compared to pungent varieties like chili peppers (Stewart Jr et al., 2005). We searched for the *Pun1* (*CA02G19260*) gene in our database and observed that it was expressed mainly in fruit and pericarp tissues, consistent with the site of capsaicin accumulation in the plant (Figure 6A). Next, we obtained the list of top co-expressed genes for *Pun1* and compared it against a list of putative capsaicin biosynthetic pathway genes previously obtained by isolation from *Capsicum annuum* fruit tissue at specific developmental stages (Mazourek et al., 2009; Zhang et al., 2016). This comparison yielded an overlap of 13.7% (7 genes), which we examined individually for their function and organ expression (Figure 6B). We observed that all the genes were predominantly found in the pericarp tissue, consistent with the site of capsaicin accumulation (Figure 6D). We then obtained a putative map of the capsaicin biosynthetic pathway (Zhang et al., 2016) and attempted to map our gene list to it. Surprisingly, we observed that most of the genes mapped to the pyruvate-derived arm of the pathway, suggesting that *Pun1* is primarily co-regulated with this arm of capsaicin biosynthesis (Figure 6C).

**Figure 6:**
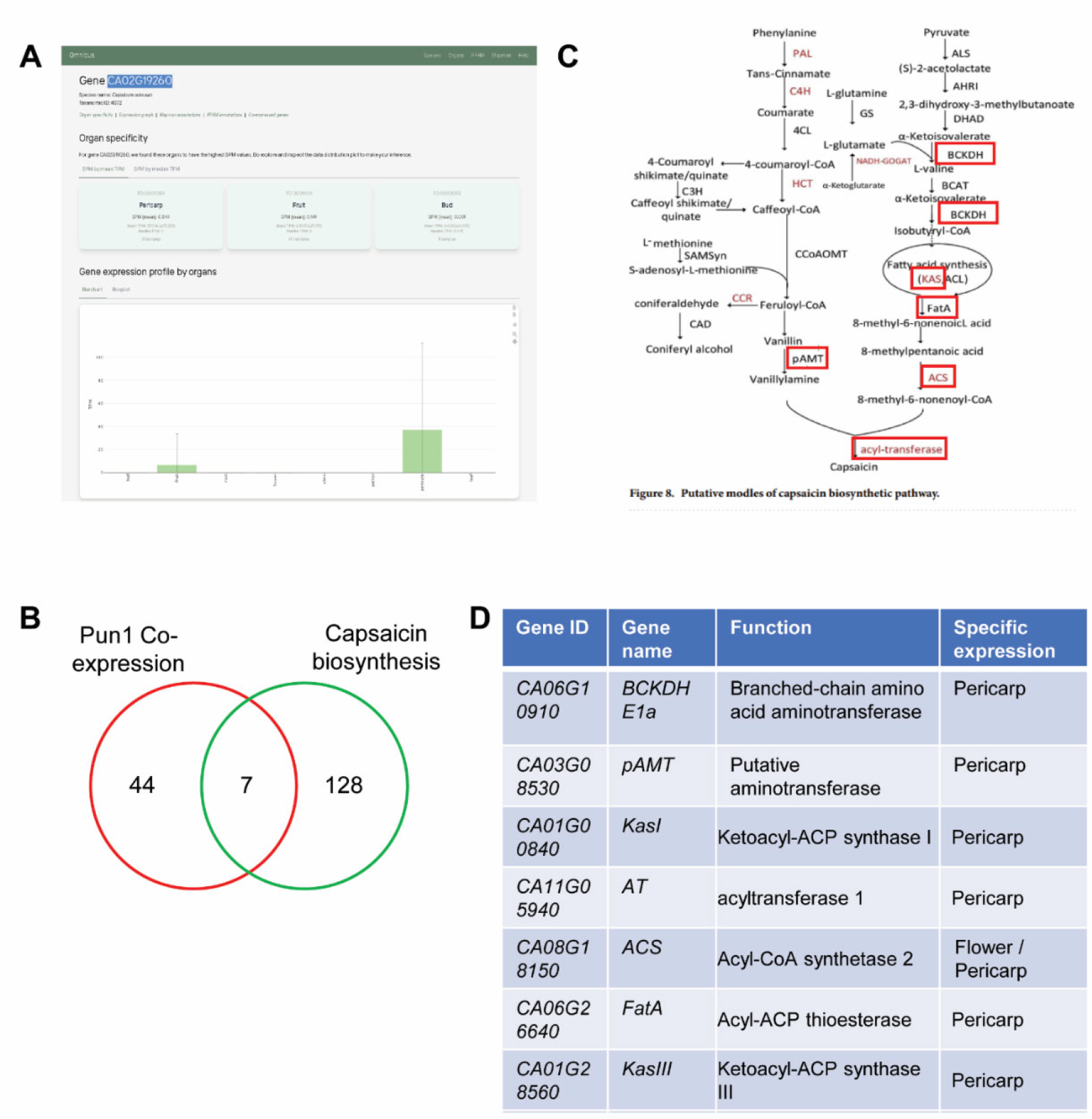
Example use case of *Pun1* (*CA02G19260*) from *Capsicum annum*, an important enzymatic component of capsaicin biosynthesis. A) Gene expression profile by organs shows its highly specific expression in fruit and pericarp, which is also the site of capsaicin accumulation. B) Comparison of top co-expressed genes with known capsaicin biosynthetic pathway components yields an overlap of 13.7% (7 genes). C) Mapping of genes to proposed capsaicin biosynthetic pathway based on putative gene function (red boxes). D) Specific expression of the seven genes obtained from PEO.

Finally, we also provide several other options for general gene and pathway discovery using either the organ, Pfam or Mapman specific annotation modes. For instance, by selecting the ‘Organs’ tab, we are directed to a page listing all specific organ / tissue types sorted by Plant Ontology (PO) number. Selecting the PO number (PO:0009005) corresponding to root directs us to a page containing genes specific to the root, sorted by SPM values (Supplemental Figure S4). Here, we can select the species of interest, in this case, *Solanum lycopersicum*, and ask PEO to provide a heatmap of the top 50 genes. As observed, the heatmap displays a set of genes showing strong enrichment in root tissue (Supplemental Figure S4).

Next, we show an example whereby Pfam IDs are used as a starting point by selecting the Pfam tab. Here we chose Pfam ID (PF00182) which corresponds to Chitinase class I, which is important for defense against fungal pathogens. We are directed to a page which displays a list of genes sorted by species that are associated with this Pfam ID (Supplemental Figure S5). Selecting *Physcomitrella patens* as our species of interest, we can ask PEO to provide a heat map of gene expression in various tissues. We observe that different genes are associated with different tissue types, providing hypothesis generation capabilities for the investigation of these genes in specific tissues.

Lastly, we also show how Mapman bins can be used as a starting point by selecting the Mapman tab. We select the Mapman bincode (12.1.1.1 Chromatin organisation.chromatin structure. DNA wrapping.histone *(H2A)) as an example. PEO directs us to a page where we are provided with a list of genes associated with this bincode sorted by species. In this case, we generate a heatmap of the species Oryza sativa, where we observe that the majority of the genes are associated with meristematic or reproductive tissue (Supplemental Figure 6). Again, this simple analysis can provide useful insights into gene function and any tissue specific properties it may possess as a starting point for experimental investigations.

## Conclusion

We present here a fully annotated Gene Expression Omnibus of 103 plant species, which provides information on tissue specificity and gene function information of the relevant gene of interest. PEO also automatically provides a list of top-coexpressed genes for better elucidation of gene functions and associated pathways. We provided two case studies of how PEO can be used to study genes and pathways of interest and to narrow down large putative gene lists based on information on tissue specificity, annotated Pfam and Mapman functions, and comparison using complementary datasets obtained from other analyses. These analyses are amenable to any species present in PEO and allow researchers to build a genetic model of a pathway of interest quickly. As we expand the number of species available in PEO, we will be able to more effectively empower researchers working on non-model organisms and more quickly progress the pace of translational research.

## Acknowledgments

We would like to thank Riccardo Delli Ponti for the initial help in collecting the gene expression data. M.M. is supported by the Singaporean Food Agency grant SFS_RND_SUFP_001_05 and E.K. is supported by Ministry of Education Tier 3 Grant ‘From Tough Pollen to Soft Matter’.

