## Supplemental Table S1-2 for "PEO: Plant Expression Omnibus - a comparative transcriptomic database for 103 Archaeplastida"

| Data | Content | Prepared by |
|---|---|---|
| Species list | Includes list of 103 species, with the sources of their CDS. | Manual curation by Mutwil. |
| TPM matrices | Quantifies gene expression of each gene in each sample. Each species is represented by a 2D TPM matrix with genes along the rows and samples along the columns. | TPM matrices were processed and compiled from other projects (Goh & Mutwil, 2021; Julca et al., 2021). Villanueva and Mutwil performed quality control. |
| Organ annotations | Assignment of each RNA-seq sample to a PO term and name. | Manual curation and annotation by Villanueva et. al. (2022, unpublished work). |
| Mapman annotations | Assignment of each gene to at least one Mapman bin label and name. | Performed in this study by Goh. |
| Pfam annotations (Interpro) | Assignment of each gene to Pfam domains, which in turn have a name and their associated GO terms. | Manual curation and annotation by Villanueva et. al. (2022, unpublished work). |
| PEP files | Protein sequence files, required for building the diamond reference db. Diamond (Buchfink et al., 2015) is an alternative to BLAST for fast protein sequence search. | Raw pep files obtained by Villanueva, re-processed by Goh. Diamond protein sequence database was processed by Goh. |

**Supplemental Table S1:** Raw data collected for PEO

| File type | Columns required | Number of files | Constraints |
|---|---|---|---|
| Species list | Taxonomic ID, scientific name, CDS source, CDS url | 1 globally | - |
| TPM matrices | Gene IDs as row index, sample labels as column headers, TPM as values | 1 per species | Species must be specified in species list |
| Gene annotations | Gene annotation label, name any other fields (flexible) | 1 per annotation type globally (currently 2, for Mapman and Interpro) | - |
| Gene annotations assignment | Gene ID, Gene annotation label | 1 per species, per annotation type | Genes must exist in TPM matrices, gene annotation must exist in globally defined file |
| Sample annotations | JSON mapping of sample annotation label (PO terms) to the name of each PO term | 1 per sample annotation type globally (1 for PO currently) | - |
| Sample annotations assignment | Sample label, Sample annotation label (PO term) | 1 per species, per sample annotation type | Samples must exist in TPM matrices, sample annotation must exist in globally defined file |

**Supplemental Table S2:** Processed data files uploaded to PEO
