## Supplementary figures and images for "PEO: Plant Expression Omnibus - a comparative transcriptomic database for 103 Archaeplastida"

### Figure S1

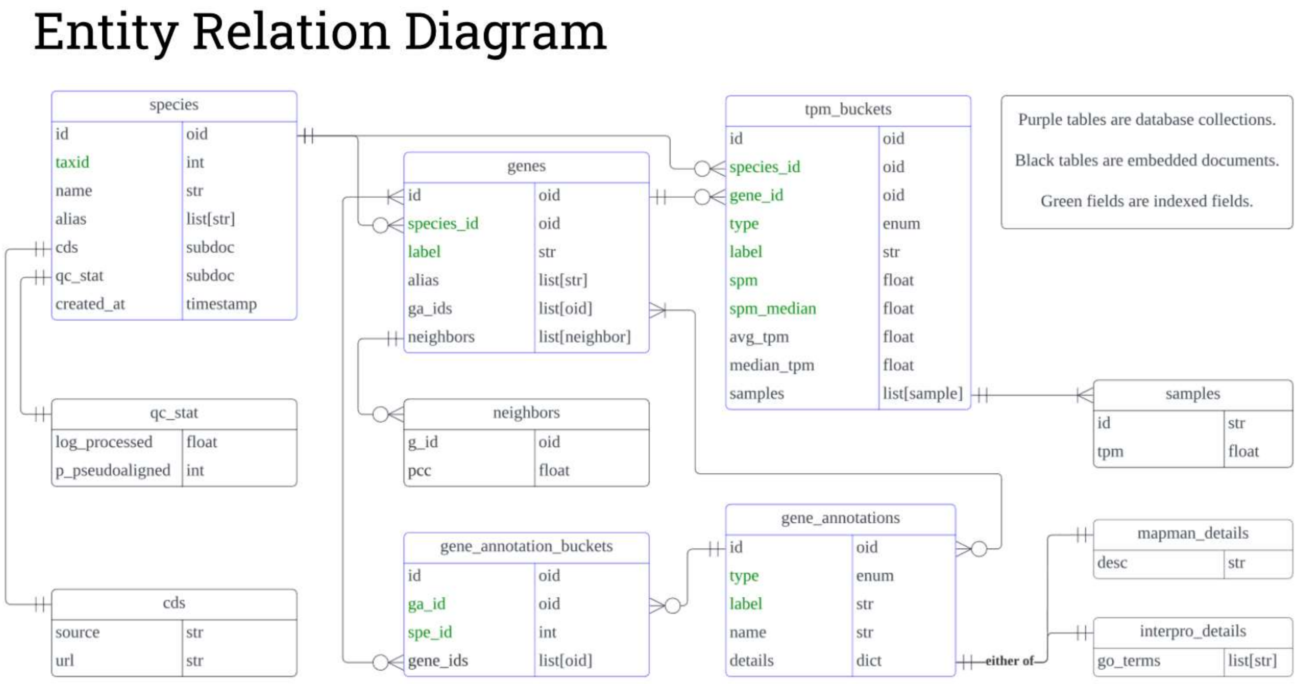


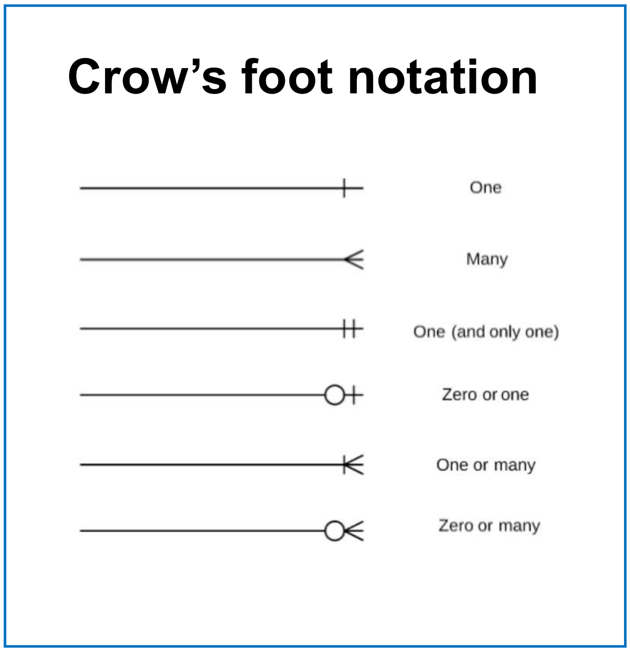


**Supplemental Figure S1:** Detailed architectural design of PEO web-tool
