## Supplementary material for "PEO: Plant Expression Omnibus - a comparative transcriptomic database for 103 Archaeplastida": Figure S3-6

#### Example: Input protein sequence (AT2G16910)

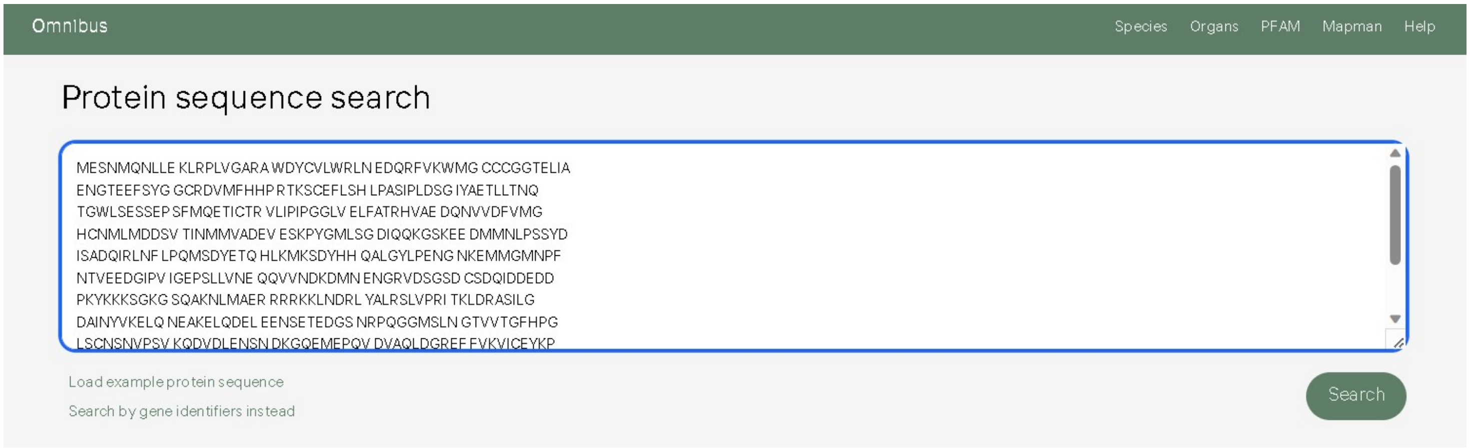


#### Output: Hits sortable by species and % identity

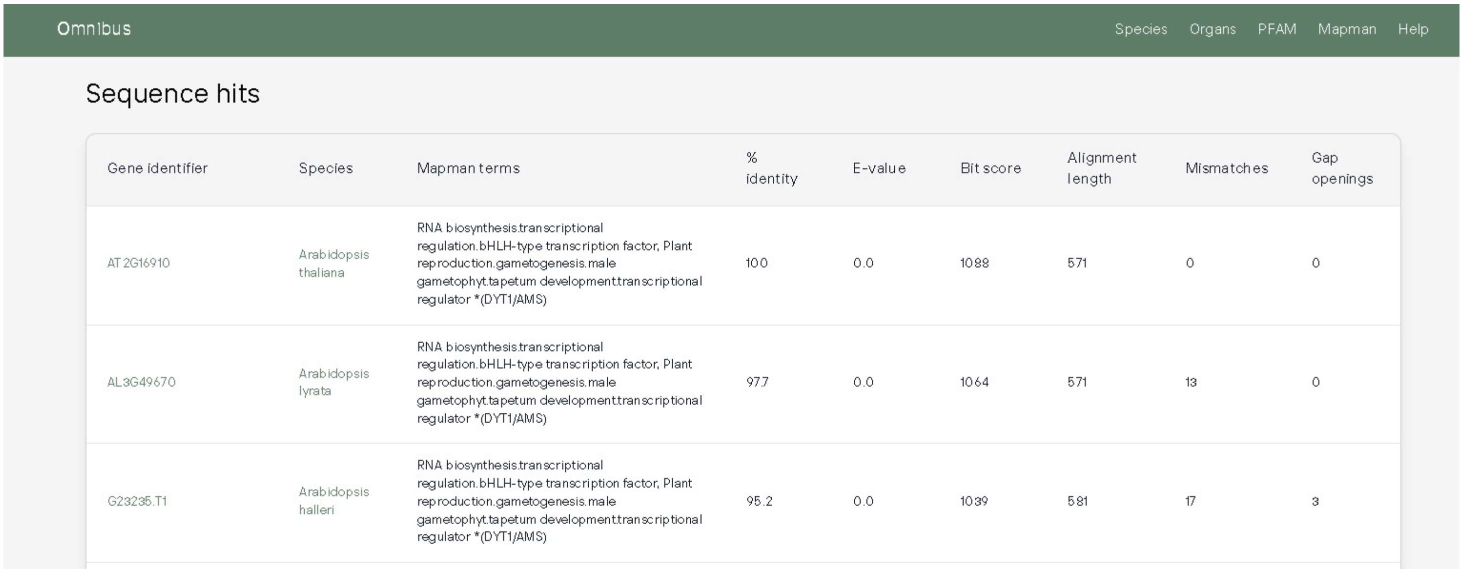


**Supplemental Figure S3:** Example search output for protein sequence. Search output retrieves and sorts candidate orthologous genes from other species available in the database.

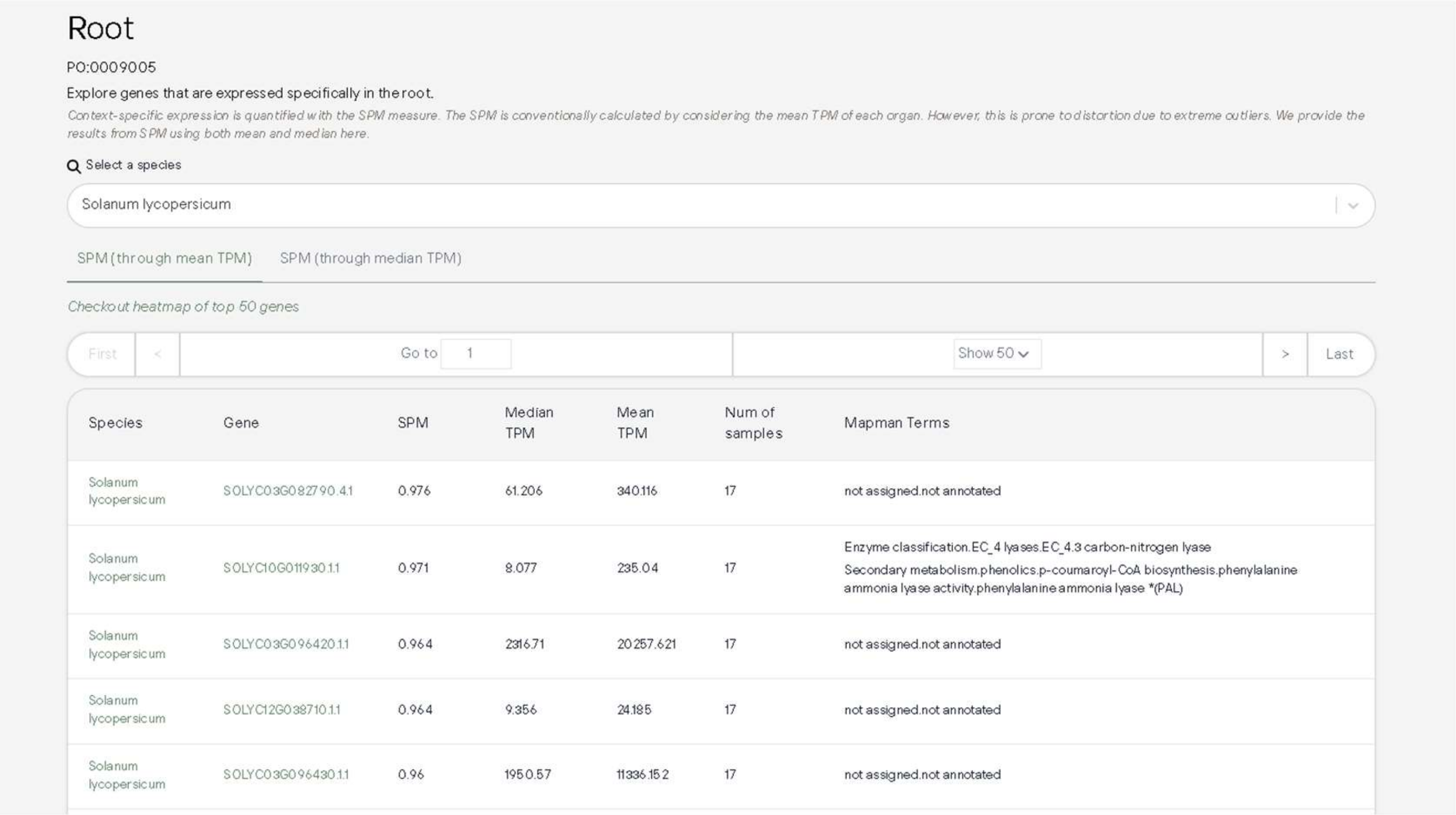

### Root

PO:0009005

Explore genes that are expressed specifically in the root.

*Context-specific expression is quantified with the SPM measure. The SPM is conventionally calculated by considering the mean TPM of each organ. However, this is prone to distortion due to extreme outliers. We provide the results from SPM using both mean and median here.*

Select a species

Solanum lycopersicum

SPM (through mean TPM)   SPM (through median TPM)

*Checkout heatmap of top 50 genes*

First   <   Go to 1   Show 50   >   Last

| Species | Gene | SPM | Median TPM | Mean TPM | Num of samples | Mapman Terms |
|---|---|---|---|---|---|---|
| Solanum lycopersicum | SOLYC03G082790.4.1 | 0.976 | 61.206 | 340.116 | 17 | not assigned.not annotated |
| Solanum lycopersicum | SOLYC10G011930.1.1 | 0.971 | 8.077 | 235.04 | 17 | Enzyme classification.EC_4 lyases.EC_4.3 carbon-nitrogen lyase<br>Secondary metabolism.phenolics.p-coumaroyl-CoA biosynthesis.phenylalanine ammonia lyase activity.phenylalanine ammonia lyase *(PAL) |
| Solanum lycopersicum | SOLYC03G096420.1.1 | 0.964 | 2316.71 | 20257.621 | 17 | not assigned.not annotated |
| Solanum lycopersicum | SOLYC12G038710.1.1 | 0.964 | 9.356 | 24.185 | 17 | not assigned.not annotated |
| Solanum lycopersicum | SOLYC03G096430.1.1 | 0.96 | 1950.57 | 11336.152 | 17 | not assigned.not annotated |

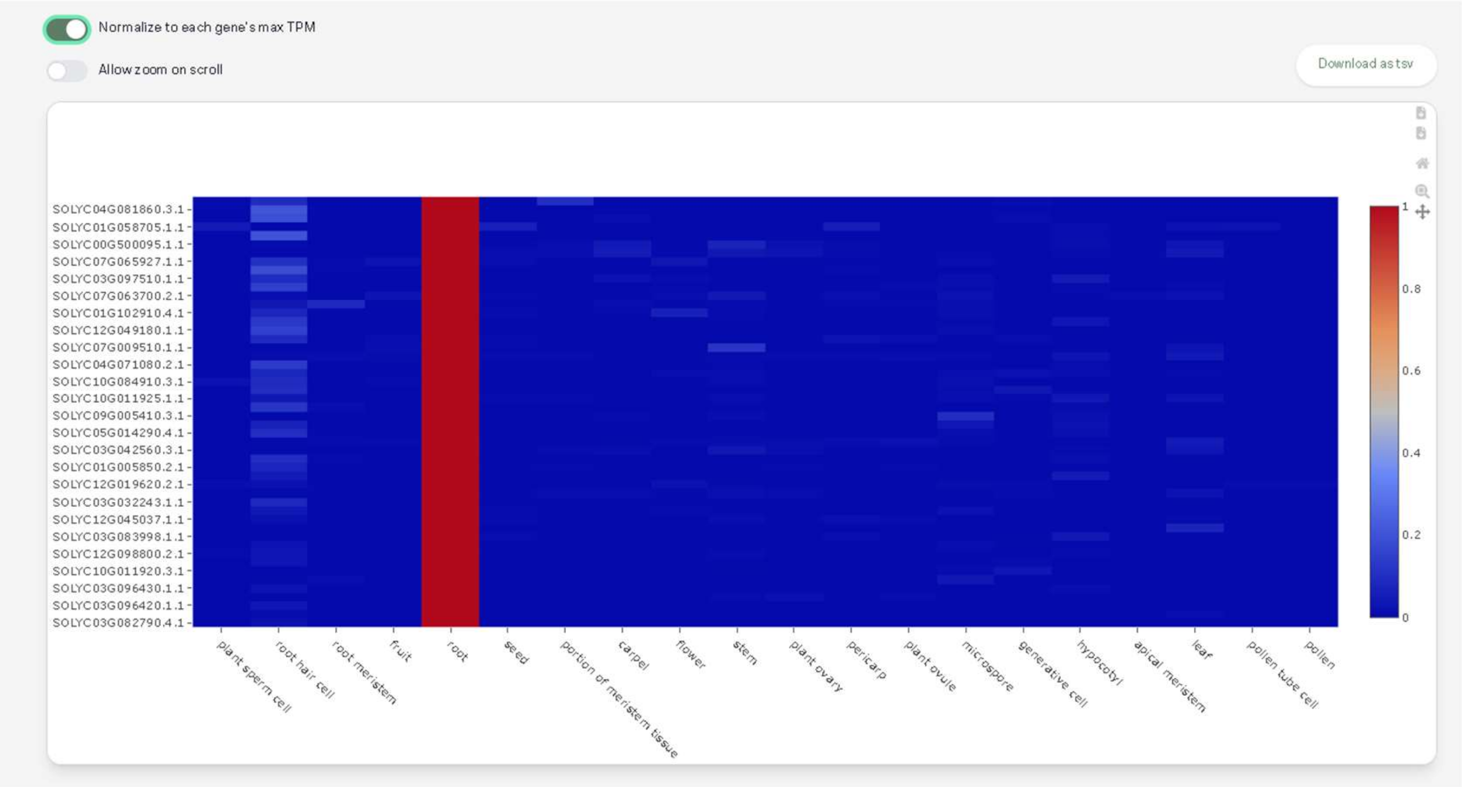


**Supplemental Figure S4:** Heatmap of organ specific genes (Top 50 genes). Heatmap of top 50 root-specific genes sorted by organ type in Solanum lycopersicum. Scale represents gene expression normalised to max TPM of individual gene.

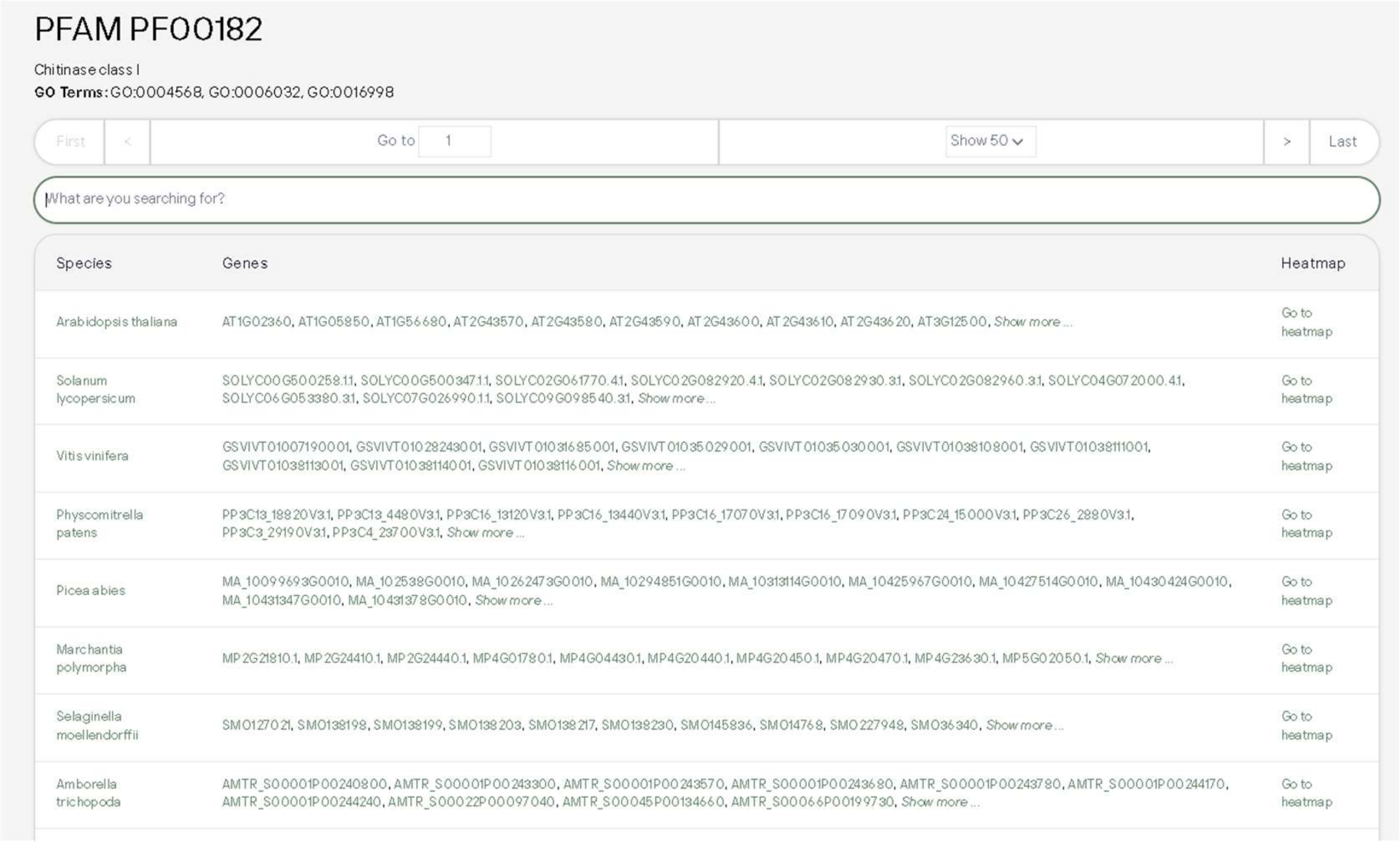

PFAM PF00182

Chitinase class I

**GO Terms:** GO:0004568, GO:0006032, GO:0016998

First < Go to 1 Show 50 > Last

What are you searching for?

| Species | Genes | Heatmap |
|---|---|---|
| Arabidopsis thaliana | AT1G02360, AT1G05850, AT1G56680, AT2G43570, AT2G43580, AT2G43590, AT2G43600, AT2G43610, AT2G43620, AT3G12500, *Show more ...* | Go to heatmap |
| Solanum lycopersicum | SOLYC00G500258.1.1, SOLYC00G500347.1.1, SOLYC02G061770.4.1, SOLYC02G082920.4.1, SOLYC02G082930.3.1, SOLYC02G082960.3.1, SOLYC04G072000.4.1, SOLYC06G053380.3.1, SOLYC07G026990.1.1, SOLYC09G098540.3.1, *Show more ...* | Go to heatmap |
| Vitis vinifera | GSVIVT01007190001, GSVIVT01028243001, GSVIVT01031685001, GSVIVT01035029001, GSVIVT01035030001, GSVIVT01038108001, GSVIVT01038111001, GSVIVT01038113001, GSVIVT01038114001, GSVIVT01038116001, *Show more ...* | Go to heatmap |
| Physcomitrella patens | PP3C13_18820V3.1, PP3C13_4480V3.1, PP3C16_13120V3.1, PP3C16_13440V3.1, PP3C16_17070V3.1, PP3C16_17090V3.1, PP3C24_15000V3.1, PP3C26_2880V3.1, PP3C3_29190V3.1, PP3C4_23700V3.1, *Show more ...* | Go to heatmap |
| Picea abies | MA_10099693G0010, MA_102538G0010, MA_10262473G0010, MA_10294851G0010, MA_10313114G0010, MA_10425967G0010, MA_10427514G0010, MA_10430424G0010, MA_10431347G0010, MA_10431378G0010, *Show more ...* | Go to heatmap |
| Marchantia polymorpha | MP2G21810.1, MP2G24410.1, MP2G24440.1, MP4G01780.1, MP4G04430.1, MP4G20440.1, MP4G20450.1, MP4G20470.1, MP4G23630.1, MP5G02050.1, *Show more ...* | Go to heatmap |
| Selaginella moellendorffii | SM0127021, SM0138198, SM0138199, SM0138203, SM0138217, SM0138230, SM0145836, SM014768, SM0227948, SM036340, *Show more ...* | Go to heatmap |
| Amborella trichopoda | AMTR_S00001P00240800, AMTR_S00001P00243300, AMTR_S00001P00243570, AMTR_S00001P00243680, AMTR_S00001P00243780, AMTR_S00001P00244170, AMTR_S00001P00244240, AMTR_S00022P00097040, AMTR_S00045P00134660, AMTR_S00066P00199730, *Show more ...* | Go to heatmap |

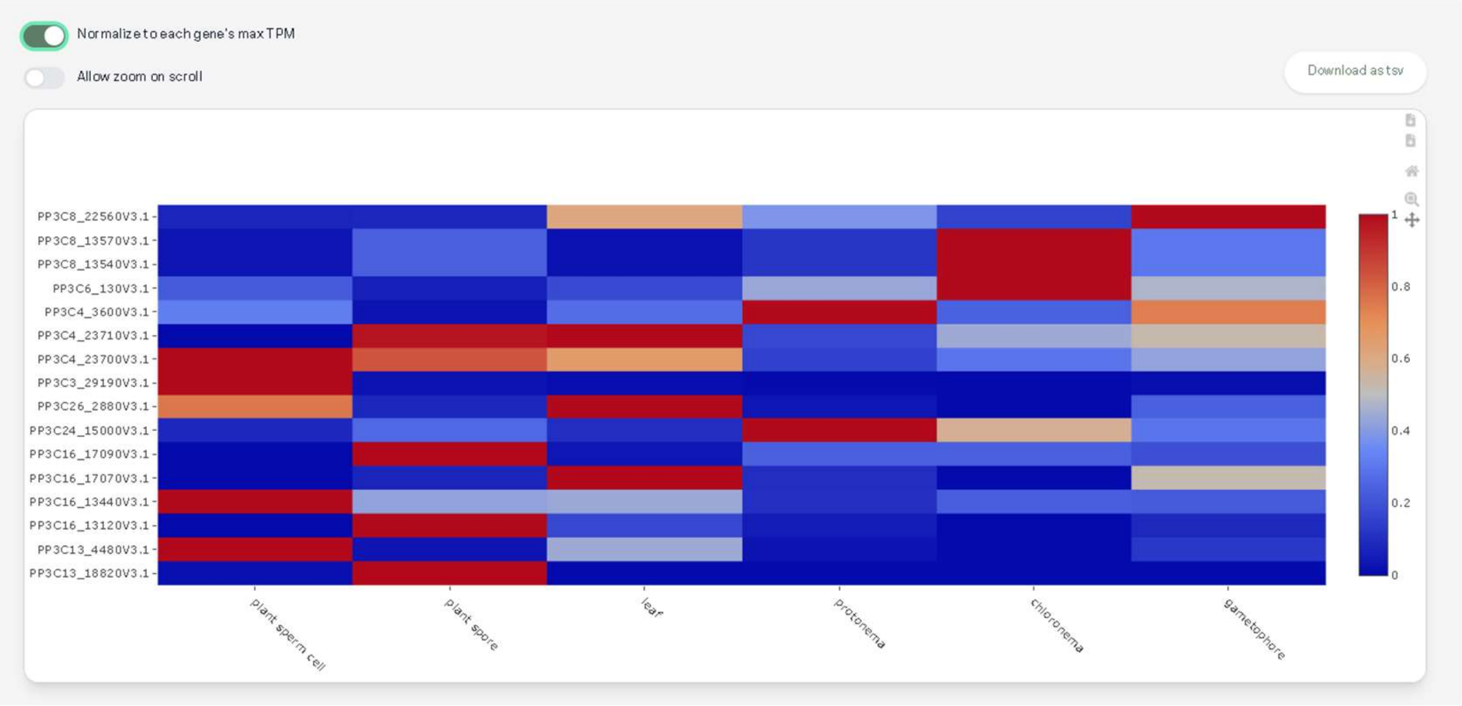


**Supplemental Figure S5:** Heatmap of genes classified by Pfam ID. Heatmap of genes classed under Pfam ID (PF00182, Chitinase Class I) sorted by organ type in Physcomitrella patens. Scale represents gene expression normalised to max TPM of individual gene.

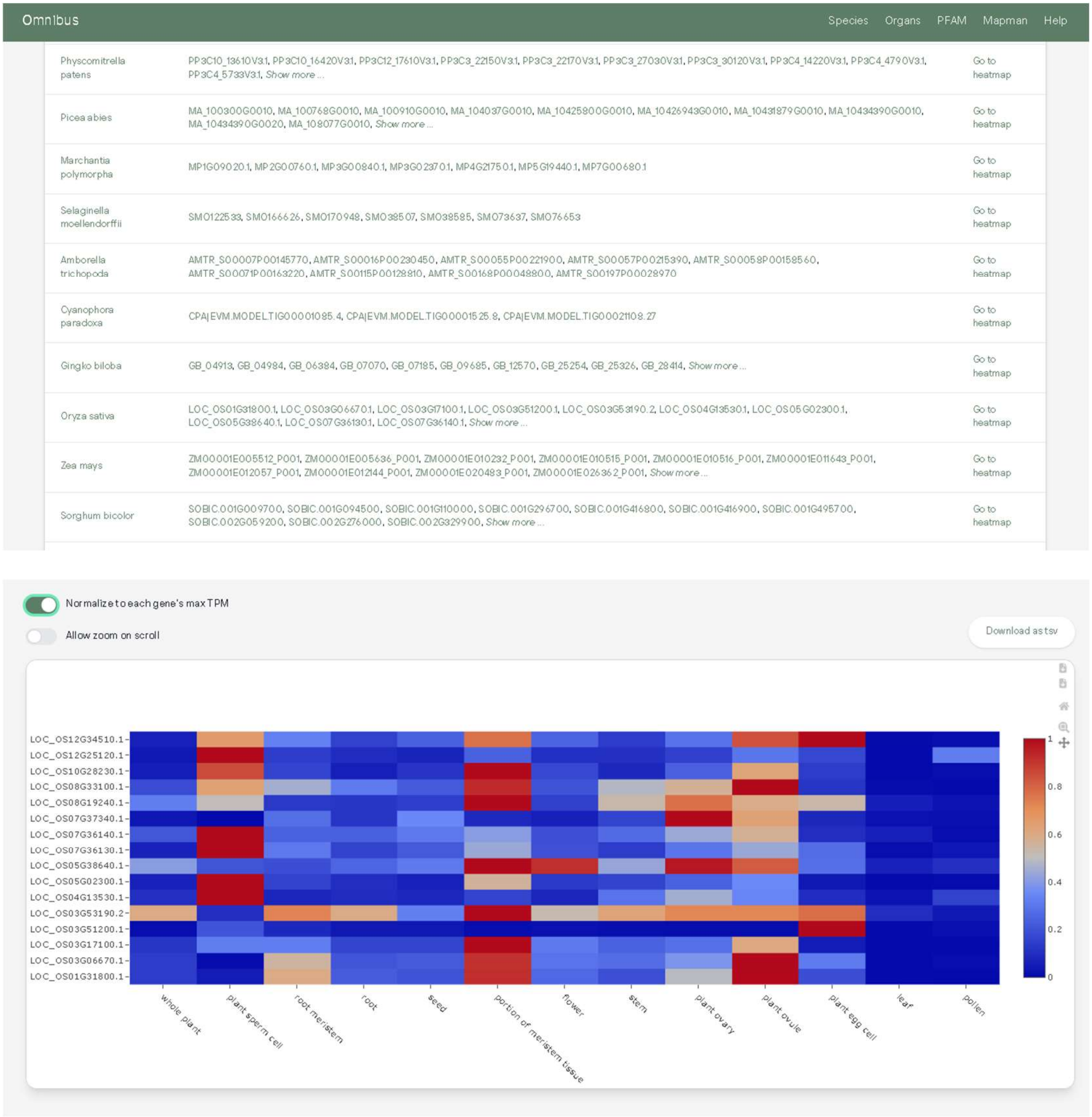


**Supplemental Figure S6:** Heatmap of genes classified by Mapman bincode. Heatmap of genes classed under Mapman bincode (12.1.1.1 Chromatin organisation.chromatin structure. DNA wrapping.histone *(H2A)) sorted by organ type in Oryza sativa. Scale represents gene expression normalised to max TPM of individual gene.
